# Tritrophic interactions involving a dioecious fig tree, its fig pollinating wasp and fig nematodes

**DOI:** 10.1101/736652

**Authors:** J. Jauharlina, Hartati Oktarina, Rina Sriwati, Natsumi Kanzaki, Rupert J. Quinnell, Stephen G. Compton

## Abstract

Many species of fig trees (*Ficus* spp., Moraceae) have nematodes that develop inside their inflorescences (figs). Nematodes are carried into young figs by females of the trees’ host-specific pollinating fig wasps (Agaonidae) that enter the figs to lay their eggs. The majority of Asian fig trees are functionally dioecious. Pollinators that enter figs on female trees cannot reproduce and offspring of any nematodes they carry will also be trapped inside. The biology of the nematodes is diverse, but poorly understood. We contrasted the development of nematodes carried by the pollinating fig wasp *Ceratosolen solmsi marchali* into figs on male and female trees of *Ficus hispida* in Sumatra, Indonesia. Figs were sampled from both male and female trees over a six-month period, with the nematodes extracted to record their development of their populations inside the figs. Populations of three species of nematodes developed routinely inside figs of both sexes: *Caenorhabditis* sp. (Rhabditidae), *Ficophagus cf. centerae* and *Martininema baculum* (both Aphelenchoididae). This is the first record of a *Caenorhabditi*s sp. associated with *F. hispida*. Mean numbers of nematodes reached around 120-140 in both male and female figs. These peak population sizes coincided with the emergence of the new generation of adult fig wasps in male fig trees. We conclude that figs on female trees can support development and reproduction of some nematode species, but the absence of vectors means that their populations cannot persist beyond the lifetime of a single fig. Just like their fig wasp vectors, the nematodes cannot avoid this routine source of mortality.

## Introduction

Fig trees (*Ficus* spp., Moraceae) often produce large crops of figs all year around. This has resulted in many species of vertebrates feeding on ripe figs, more than recorded for any other plants [1]. Figs are also fed upon by a wide range of invertebrate species, including wasps, flies, beetles and moths, which in turn support a diverse parasitoid fauna [2]. Mites and nematodes are also found in some figs [3-5] as well as microrganisms such as fungi [6] and protistans [11].

Female pollinator fig wasps (Agaonidae) enter figs in order to lay their eggs inside the ovules that line their inner surface [7]. Their larvae develop inside the ovules, which are galled by the females at the time that eggs are laid [8]. This is in contrast to most other fig wasps (non-pollinating fig wasps-NPFW) that usually do not enter figs to oviposit, but lay their eggs into the ovules while standing on the outer surface of the figs The entry of fig wasp pollinators into figs allows them to be used as vectors for transport between figs by a variety of smaller less intrinsically mobile organisms, including microorganisms, mites and nematodes [9-11].

Fig trees display two contrasting breeding systems. Monoecious fig trees have trees where individual figs that contain both female and male flowers and the female flowers support the development of both seeds and pollinator fig wasp offspring. Dioecious fig tree species have individuals that either produce ‘male’ figs that support development of fig wasp offspring and pollen or have ‘female’ figs that reproduce via seeds [12, 13]. Adult female pollinator fig wasps that enter receptive female figs cannot reproduce and often do not re-emerge from the first fig they enter. Even if they do re-emerge, they have lost their wings and cannot fly away in search of figs on other trees. Reproduction in dioecious fig trees is maintained because of mutual mimicry between the sexes, which results in pollinator females failing to distinguish male from female figs prior to entry [14] and because once they are inside the figs the pollinators continue to behave as if they were in male figs [15]. The inability of pollinators to distinguish between male and female figs means that any animals that are transported between figs of a dioecious fig tree are routinely at risk of being taken inside a female fig, from which there will be no subsequent generation of fig wasps to act as vectors and at certain seasons most of the figs available for entry may be female. Perhaps reflecting this significant potential source of mortality, published records suggest that phoretic mites are only associated with monoecious fig tree hosts [5]. In contrast, nematodes are associated routinely with both monoecious and dioecious fig trees species [11].

Fig trees and their pollinators have a long history of mutualistic association, extending for tens of millions of years [16] and Dominican amber fossils show that pollinator fig wasps have been also transporting nematodes between figs for much of this period [17]. Today, nematodes are recorded throughout the distributional and taxonomic range of fig trees [18-21]. The nematodes in figs belong to several different families, suggesting multiple independent colonisations of figs, followed in some cases by extensive radiations [22]. One fig tree species may support several different species of nematodes and up to eight species of nematodes have been recorded from a single fig of *F. racemosa* L. in Indonesia [11]. Nematodes develop and reproduce inside the figs and offspring are ready to attach themselves again to female pollinators when the new generation of adult pollinators are ready to leave the figs [3, 10]. Female pollinators are chosen preferentially by nematodes, because males and most female NPFW do not enter the figs to oviposit so cannot act as vectors in the same way as female pollinators [23, 24]. Transfer of nematodes into figs via the ovipositors of NPFW has not been confirmed [5, 10, 25]. *Schistonchus caprifici* Gasperrini, a nematode that reproduces in male figs of *Ficus carica* L., has nonetheless been recorded as also entering NPFW females, although there is no evidence that they ever manage to enter the figs and reproduce [25]. Nematodes waiting to attach to female pollinators may be scattered around the interior of the figs, or be aggregated in male flowers [19]. The latter is a response to the active pollen collection behaviour of some female pollinators [26]. These females seek out the male flowers and move pollen from them into their pollen pockets, and collect the nematodes while doing so.

The feeding behaviour of most fig nematodes is unknown, but is clearly diverse [27, 28]. The presence of stylets is indicative of plant-feeding, but different species vary in their preferred feeding sites within the figs [28]. Nematodes use their stylets to puncture plant cells, to withdraw food and also to secrete proteins and metabolites that aid the nematode in feeding on the plants [29]. Among species that lack stylets, some feed on the decaying corpses of pollinator females, and some may also start feeding on the females before they have died [30]. Fig trees belonging to subgenus Sycomorus often have their figs partly filled by a liquid at times during their development [31] and these species appear to support particularly rich nematode faunas. Free-swimming nematodes in these figs may be predatory on other nematodes or feed on the protistans that are often present at high densities in the fig liquid [11].

Males and females of dioecious plants can differ in their attractiveness and suitability for plant-feeding invertebrates [32]. ye. Figs of both tree sexes fill with liquid in Sycomorus figs, but only male figs contain male flowers, and although the figs of both male and female plants contain female flowers, their contrasting floral development (with galled ovules or seeds, respectively) mean they offer differing resources to any nematodes that have been carried inside. In this paper, we address the following questions in relation to the nematodes associated with a dioecious fig tree, *F. hispida*, in Indonesia: (i) How many nematode species are present locally, and how abundant are they? (ii) What are their life cycles? And (iii) Are they capable of developing and reproducing inside both male and female fig trees?

## Materials and methods

### Study species and site

*Ficus hispida* is a fig trees belonging to subgenus Sycomorus and is functionally dioecious with distinct male and female individuals. The species is distributed throughout India, Nepal, Laos, Thailand, Malaysia, southern China [33], Sri Lanka, Myanmar, New Guinea, Australia, Andaman island [34], and also Indonesia [28]. *F. hispida* is a shrub or moderate-sized free standing tree up to 13 metres tall, with spreading branches. Like many other dioecious fig trees, *F. hispida* sometimes shows asynchronous fruiting within a plant, with different phases of figs found at the same time on the same tree. Pollinating wasp that enter figs from female trees will pollinate the flowers, but are unable to lay eggs because of stigma structure and because the figs contain only long-styled female flowers that prevent the wasps’ ovipositors from reaching the ovules. Therefore, mature female *F. hispida* figs contain seeds, but no pollinator offspring. The pollinating wasps that enter figs of male trees can lay eggs in the female flowers, where the fig wasp offspring develop, but no seeds are produced. Male flowers inside male figs mature at the same time as the pollinator offspring, allowing them to transport pollen to other trees. The newly emerged female pollinators disperse to find new receptive figs and start another developmental cycle if they enter figs on a male tree [7, 35, 36].

Figs of *F. hispida* are borne in long clusters at the base of the tree, as well as on the branches [7]. Fig development has been characterised by Galil and Eisikowich [38] with modifications by Valdeyron and Lloys [39] for dioecious figs. Phase A figs are pre-receptive. Pollinators (and nematodes) enter during B phase. Fig wasp offspring (male trees) and seeds (female trees) develop and mature during C phase, along with any nematodes reproducing inside the figs. In D phase the next generation of male pollinators emerge to mate with females that are still in their galls. The females then leave their galls, actively collect pollen into pollen baskets and emerge through an exit hole in the fig wall cut by the males (the start of E phase). Figs from female trees do not have a D phase, as no pollinator wasps develop there. Once the seeds in female figs are mature, the figs ripen and become attractive to seed dispersers (E-Phase). Like many other fig trees belonging to subgenus Sycomorus, C phase figs of *F. hispida* often contain noticeable amounts of liquid [40]. *C. solmsi marchali* is the only pollinator recorded from *F. hispida* in Indonesia [28], but *F. hispida* is host to different *Ceratosolen* pollinators elsewhere within its wide geographical range. Its larval development was described by [41].

The phenology and contents of *F. hispida* figs were monitored in the northern part of Sumatra Island in Aceh Province, Indonesia for six months from March to August 2018. The trees were growing along roads in mountainous areas of Leupung District, about 25 km (95° 15’ 34.92” E; 05° 22’ 55.68” N) from the provincial capital, Banda Aceh. Roadside *F. hispida* are common in the study area, usually growing in clumps of several individuals with both female and male trees present together. The region has a tropical climate that supports rainforest vegetation, with fairly constant average temperatures throughout the year and little diurnal variation. Meteorological information for the area was obtained from Blang Bintang Station, the closest Meteorological Station under the Indonesian Meteorological and Geophysical Agency, located about 30 km from the study area. There was little seasonal variation in daily temperatures during the six months of study. The average daily temperature during the study was 28.22 ± 0.09 0C (mean ± SE), with average minimum temperature was 24.18 ± 0.09 0C (mean ± SE) and maximum was 32.26 ± 0.15 (mean ± SE). Monthly minimum and maximum temperatures are quite stable, with only slight variation between the six months of study. During the six months of the study, March was the driest month with no rain at all, while May was the wettest month with 15 days of rain that accumulated 361.7 mm of precipitation.

Figs were present on both sexes of *F. hispida* more or less continuously. Asynchronous fruiting often resulted in a variety of developmental stages being present at any one time. Reasonably discrete ‘cohorts’ were nonetheless present, allowing the development of marked groups of figs to be followed. Sampling followed the complete development of each cohort. In some cases, mature D and E phase figs were present on the trees when new A-phase figs first appeared, providing opportunities for some cycling of fig wasp populations on individual male trees. Fig cohorts were sampled at weekly intervals from three male trees and three female trees (Table 1).

**Table 1.**
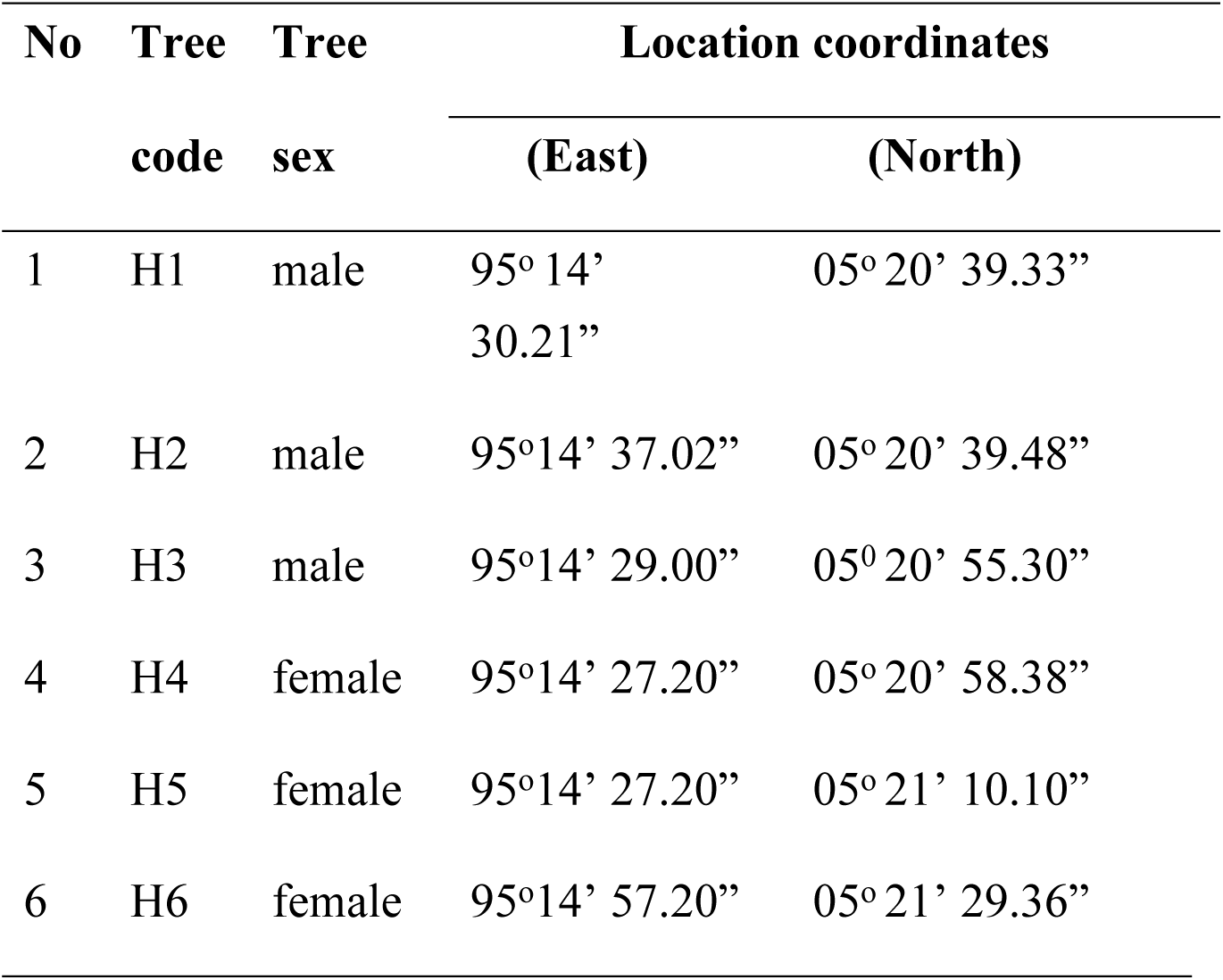
Location of the *Ficus hispida* trees sampled.

### Sampling procedures and fig extractions

The development times for each cohort were calculated from when the first A phase figs were recorded until the first E phase fig was present. Ten figs were sampled haphazardly from each cohort of each tree from A phase through to E phase (when wasp offspring had left the male figs, and female figs were soft and ready to be eaten by frugivores). The developmental stage, colour, and diameter of each fig were recorded. Later the same day, the contents of each fig, including any liquid if present, were placed individually in a Baermann extraction funnel, using a method adapted from Sriwati, Takemoto and Futai [42] as modified by Jauharlina [11]. Each fig was cut into six to eight pieces and placed onto a layer of fine fabric and immersed in 60 ml of distilled water. Water held within the funnel ensured that the fig pieces remained under water. After 24 hours, the liquid below the funnel was placed in 20 ml reaction tubes and left undisturbed for 3 hours. The upper part of the liquid was then removed without disturbing the lower liquid using a small pipette, and then discarded. Occasional checks confirmed that it did not contain nematodes. One ml extracts from the remaining 5 ml in the bottom of the tubes were placed on a one ml capacity nemacytometer glass slide (counting slide) and observed under a microscope. Any nematodes present were counted and adults were identified. Observations were repeated on the rest of the extract, giving five counts from each fig.

## Result

### Fruiting phenology of *F. hispida*

Over the six-month sampling period, a total of 430 figs were collected. The development of figs from A-phase to E-phase lasted for seven to eight weeks for male trees and for seven weeks for female trees. The figs from male trees were 1.59 ± 0.03 cm in diameter (mean ± SE, N = 30 figs) during A phase prior to pollinator entry, and reached 3.39 ± 0.06 cm in diameter (mean ± SE, N = 30 figs) at maturity. The figs from female trees were roughly the same size as male figs at similar stages of development (1.47 ± 0.07 cm, mean ± SE, N = 30, at A phase, and 3.55 ± 0.02 cm, mean ± SE, N = 30, at E phase). Figs from A, B, and C-phases were green in colour and had a hard texture with abundant white latex. When the figs developed into D-phase (male trees) and late C-phase (female trees), they became a little softer and developed a yellowish colour. The amount of latex also decreased.

B-phase was the shortest phase during the development of figs. It lasted only 2-3 days. At this stage the ostiole became a bit loose to allow the pollinating wasps to enter. The number of pollinators that entered the B-phase figs ranged from 1 - 3 wasps per fig, with an average of 1.30 ± 0.1 (Mean ± SE, N = 30 figs) on male trees and 1.27 ± 0.1 (Mean ± SE, N= 30 figs) on female trees. Average number of entries did not vary between the sexes (glmer, z = 0.11, P = 0.90). One pollinating wasp per fig was the most common occurrence in both male and female trees of *F. hispida*, representing 76.6 % and 80 % of the total figs respectively (Table 2).

**Table 2.** Frequency of pollinating fig wasps entering the B-phase figs on male and female trees of *Ficus hispida* (N=30 figs from each tree sex)

| Tree sex | Frequency of pollinating wasps entering figs (%) |  |  |
| --- | --- | --- | --- |
|  | 1 | 2 | 3 |
| Male | 76.6 | 16.7 | 6.7 |
| Female | 80.0 | 13.3 | 6.7 |

### Fig nematodes and their life cycle

A sub-sample of 185 figs (91 from male and 94 from female trees) had their contents extracted in the laboratory to examine nematode development inside the figs. Nematodes were recorded in all fig samples from male trees (100 % occupancy) and in most of the figs from female trees (91.18 % Occupancy). Three species of nematodes, a bacteria feeder, *Caenorhabditis* sp. (Rhabditidae), and two plant parasites, *Ficophagus* cf. *centerae* and *Martininema baculum* (Aphelenchoididae), were recorded, and all were found in figs from both male and female trees. *F.* cf. *centerae* and *M. baculum* were previously identified based on molecular profiles (Sriwati et al., 2017). The other species, *Caenorhabditis* sp. is typologically and phylogenetically close to another fresh fig associate, *C. inopinata* (Kanzaki et al., 2018), but its detailed taxonomic description and molecular phylogenetic status will be presented elsewhere. After being carried into the B phase figs, the nematodes were present within the central lumen throughout fig development (including the liquid if present). Old pollinating wasps that had transported the nematodes appeared to be quickly abandoned shortly after entry, and there was no indication that the corpses of the vectors continued to provide nutritional resources. 73.3% of 30 B-phase figs from male trees had pollinators with nematodes still in physical contact, ranging 1-2 nematodes per wasp with an average of 1.09 ± 0.06 nematodes per fig wasp (Mean ± SE, N = 24 pollinators). On female trees, 56.7 % of 30 B-phase figs had pollinators with nematodes, ranging 1-2 nematodes per wasp with an average of 1.18 ± 0.09 (Mean ± SE, N = 17 pollinators). All nematodes transported by the wasps into the B-phase figs were at pre-adult or juvenile stages.

The number of nematodes found inside B phase figs ranged from 3-26 per fig on male trees with an average of 10.20 ± 1.68 (Mean ± SE, N = 15 figs) and 0-20 on female trees with an average of 8.44 ± 1.57 (Mean ± SE, N = 18 figs). There was no significant difference between fig sexes in mean numbers of nematodes inside B phase figs (glmer, z = 0.459, P = 0.759) (Fig 1). At D-phase, when the new generation of fig wasps were ready to leave male figs, immature nematodes attached themselves to the female wasps and were carried out of the figs. The life cycle of fig nematodes transferred by pollinating wasps into figs of *F. hispida* trees is summarized in Fig 2.

**Fig 1.**
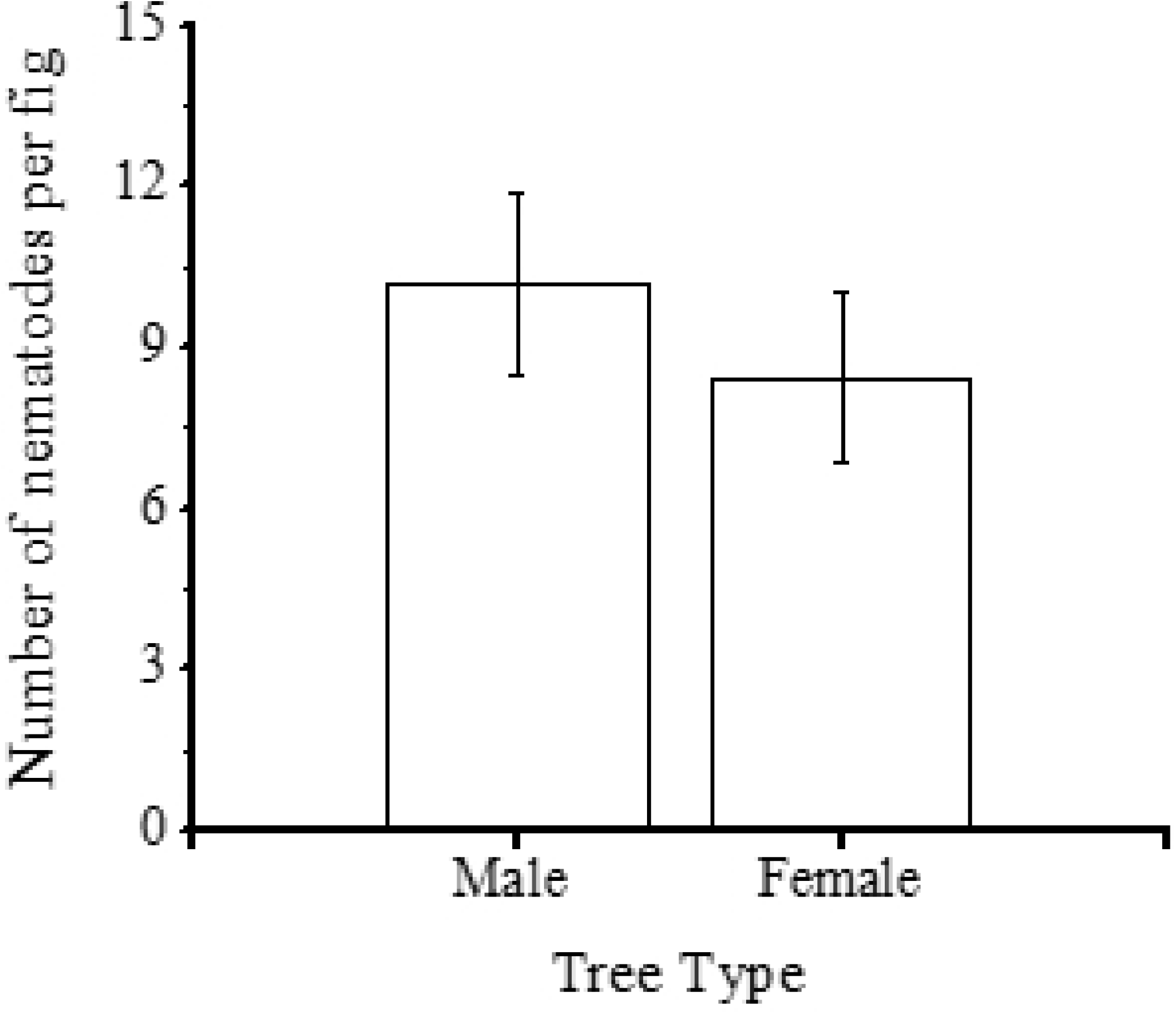
The number of nematodes found inside B-phase figs on male and female trees of *Ficus hispida*, shortly after the death of fig wasp vectors. (Mean ± SE, N = 15 figs for male trees, N = 18 figs for female trees).

**Fig 2.**
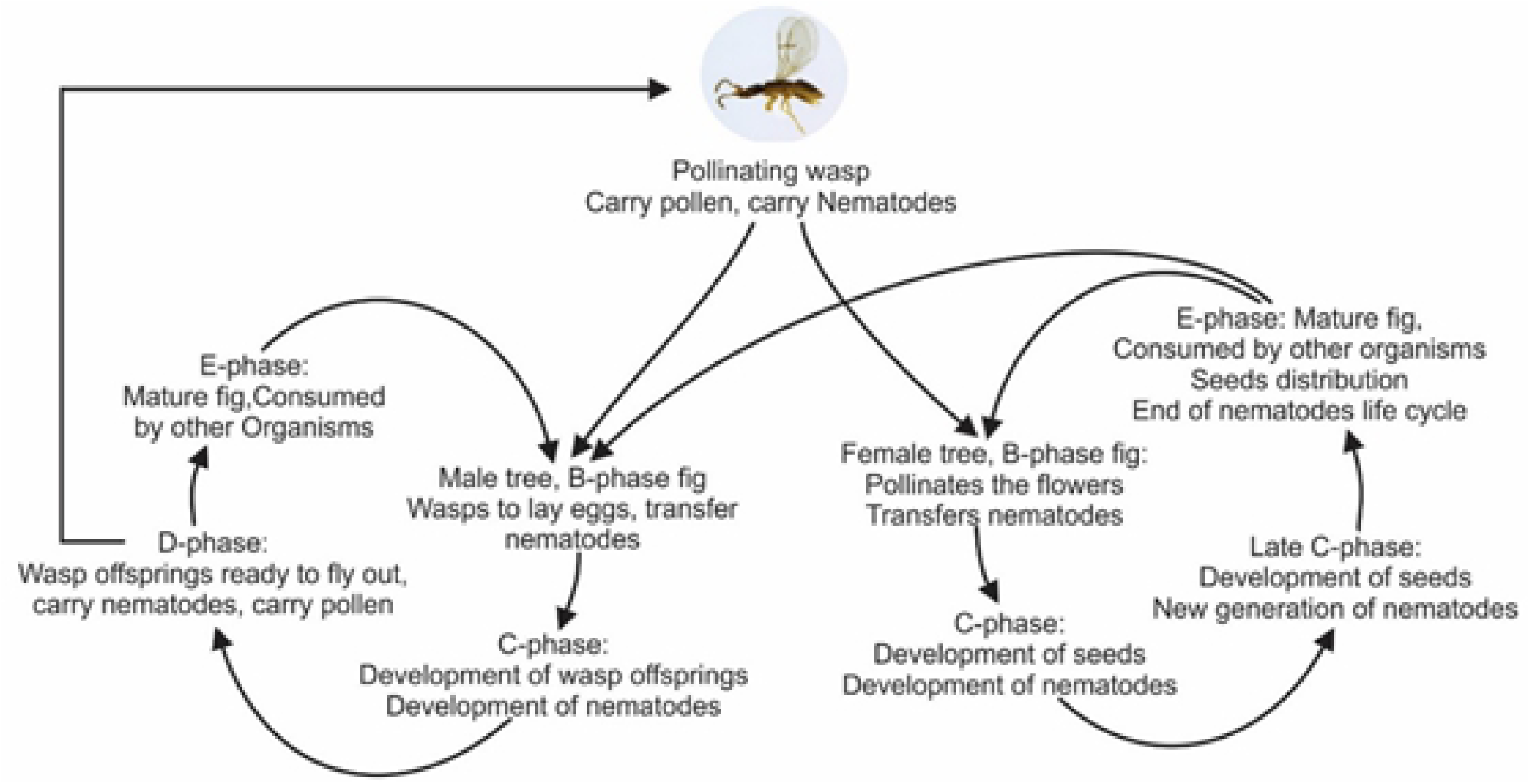
Life cycles of nematodes associated with pollinating wasps on male and female dioecious fig trees of *Ficus hispida* based on routine observation on figs morphology and extract.

All three nematode species produced offspring inside both figs from male and female trees. In male trees, peak populations of nematodes were in D-phase figs, when the new generation of wasps was ready to leave (Fig 3). In female trees, where wasps could not reproduce, all the nematodes that developed inside the figs had no means of dispersal. Peak nematode populations in female figs were at late C-phase (equivalent in timing to D-phase in male figs) and the time when they would have been seeking out fig wasp vectors, if any had been present (Fig 4). The peak population sizes in figs of both male and female trees were not significantly different from each other (glmer, z = −0.837, P = 0.402) showing that nematodes could reproduce successfully in figs of female trees of *F. hispida.*

**Fig 3.**
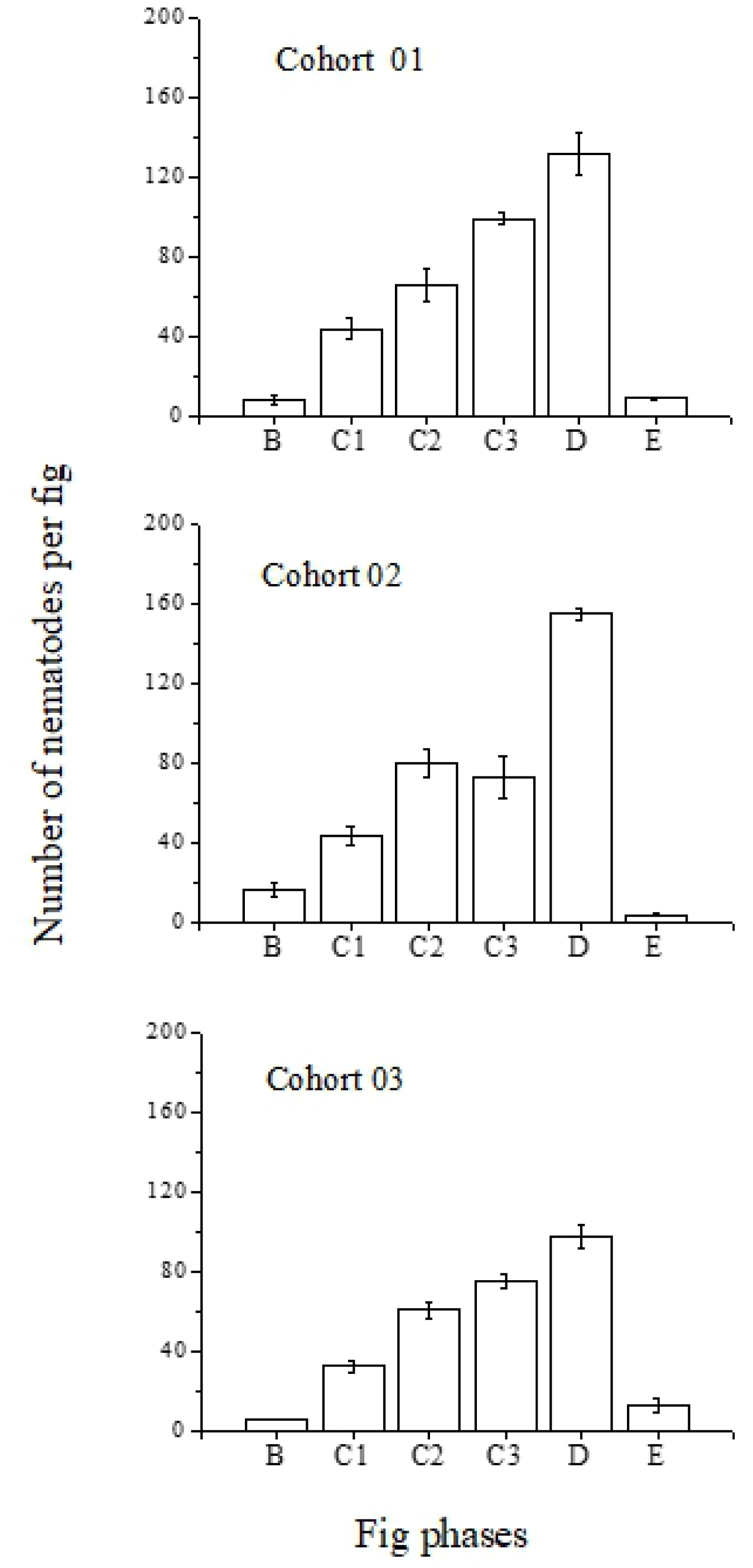
Nematode populations per fig (all species and stages) during fig development on male trees of *F. hispida*. Counts were obtained from extractions of whole figs (Mean ± SE, N = 5-6 figs for each phase, error bars represent standard errors of the means). Fig phases follow the terminology of Galil & Eisikowich [38].

**Fig 4.**
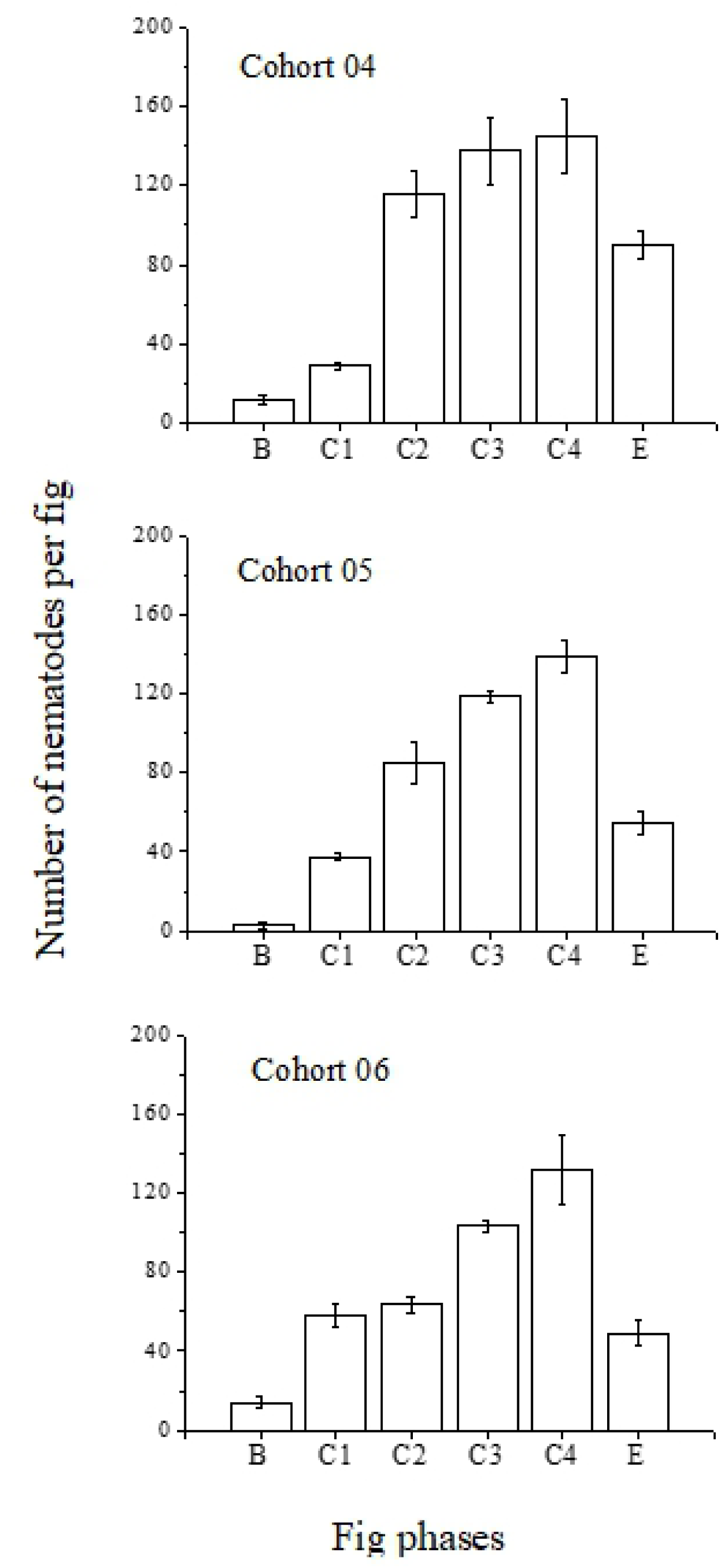
Nematode populations per fig (all species and stages) during fig development on female trees of *F. hispida*. Counts were obtained from extractions of whole figs (Mean ± SE, N = 5-8 figs for each phase, error bars represent standard errors of the means). Figs from female trees do not have a D-phase as there is no fig wasps develop inside the figs. Phases follow the terminology of Galil & Eisikowich [38].

Male fig wasps emerge before their females and seek out galls that contain the females for mating. Nematodes were seen on these males. Examination of the new generation of adult female fig wasps in early D phase figs, before holes had been chewed into the walls of their galls by the males, found no nematodes were present. This changed once the male pollinators chewed holes into the galls in order to mate, and the nematodes could gain entry. Similarly, there were no nematodes recorded from inside the anthers of male flowers. Judging from the presence of adult nematodes and immatures throughout all but the earliest phases of fig development, in both male and female figs, it is likely that both male and female figs supported more than one generation of each of the three nematode species during the time taken for one generation of fig wasps to develop.

## Discussion

Mature individuals of *F. hispida* in equatorial North Sumatra fruited almost continuously, with one fig cohort merging into another all year round. On male trees this sometimes resulted in sufficiently unsynchronized fruiting for fig wasp populations to cycle on one male tree, as seen in some other dioecious fig tree species [43, 44]. Asynchronous fruiting should reduce pollinator mortalities associated with flight between trees [41], and thereby increase the trees’ ability to maintain local populations of their pollinators and the nematodes they carry [44]. It may also increase the likelihood of pollinator females entering male figs, and increase the numbers of females entering individual male figs, both of which will be advantageous for any nematodes they are carrying.

Two of the nematode species found in this study were the same species as those described earlier from the same area [45)] Previously the two nematodes species from family Aphelenchoididae were morphologically identified as *Schistonchus centerae* and *S. guangzhouensis* [11], however further molecular identification showed that these two species were *Ficophagus* cf. *centerae* and *Martininema baculum* [45]. Aphelenchoididae species are known to be phytophagous and feed on flowers inside the figs. They have a stylet, a stomatal structure used to feed on plant tissues [20, 45]. During C phase, many nematodes were also seen swimming in those figs where liquid was present. Adults and juveniles of the two aphelenchoidid species from *F. hispida* sought out only adult female pollinators, which became available to the nematodes after pollinator males had chewed holes into the females’ galls for mating. A New World Schistonchus sp. associated with F. laevigata contacts adult pollinators in the same way [27].

Rhabditid nematodes have been recorded previously from figs of F. septica in Taiwan [46]. Recently, the species *Caenorhabditis inopinata* has been observed inside the figs of *F. septica* in Japan [47, 48] and the presence of Caenorhabditis nematodes in *F. hispida* figs is a new record. Its ecology in the figs of *F. hispida* is still unclear, but it was transferred between figs exclusively as juvenile dauer larvae. Other *Caenorhabditis* species are colonizers of nutrient and microorganism-rich organic material [9, 50]. The *Caenorhabditis* species from *F. hispida* appears not to be facultatively necromenic (feeding on the vector’s cadaver after the fig wasp dies) and given that there is more than one generation within the figs, other food sources are clearly used. Protistans are routinely present within the lumen of the figs and they are one potential food source, as are the larvae of the aphelenchoidid species.

The life cycles of the nematodes found in *F. hispida* were similar to those reported earlier in other fig tree species [5, 11, 51-53], but this study has revealed that nematodes can develop and reproduce inside figs from female trees of *F. hispida* despite the absent of fig wasp offspring. Studies of nematodes in dioecious fig trees have focused on male trees, from which a new generation of fig wasps can develop [20, 35]. Nematodes developed and reproduced within figs on female trees in a similar way, but they perished once the figs were mature and either eaten by vertebrates or fell to the ground. Resources absent from figs on female trees (galled ovules and male flowers) were clearly not required by the nematodes, but both pollinating fig wasps and the nematodes they carry are frequent victims of their host plant’s reproductive system.

Despite the high occupancy rates and large numbers of nematodes within the figs of *F. hispida* they had no obvious effect on the development of pollinating wasps inside the galls on male trees and the development of seeds on female trees. In male figs, the reproductive success of the pollinators is important for the survival of future generations of nematodes because reduced pollinator reproductive success translates into fewer vectors for the later generations of nematodes [30]. Even among the nematode species that feed on dead or dying pollinators after they enter new figs there is little evidence that they cause significant harm to the living fig wasps [54]. This apparent absence of a negative impact on the pollinators is advantageous to the nematodes, and they contribute to the exceptional biodiversity centered on figs without impinging on the core mutualism on which that diversity depends.

## Acknowledgements

The authors would like to thank for the help provided by the field work team: Yusmaini, Mardiana, and Ikram Taufik. This study was funded by Syiah Kuala University, Ministry of Research, Technology and Higher Education Indonesia, contract number: 288/UN11/SP/PNPB//2018.

